# Distinguishing “self” from “other” in a dynamic synchronization task with an adaptive virtual partner

**DOI:** 10.1101/625061

**Authors:** Merle T. Fairhurst, Petr Janata, Peter E. Keller

## Abstract

For precise interpersonal coordination, some degree of merging a sense of self with other is required. In group music making, one may want to be in “sync” with one’s ensemble and, if playing a similar instrument, one can assume a degree of temporal and acoustic overlap. However, to what extent is self-other merging optimal? An incorrect balance of segregation and integration of self and other information would result in a lack of interpersonal cohesion or a disruption of self-agency. Using an interactive finger-tapping task with a virtual partner and functional MRI, we explored neural differences between self-other merging and distinction. Varying both the level of adaptivity of a virtual partner and the quality of self-related auditory feedback, we show that the predictability of the other and availability of distinguishable, self-related information improve performance and demonstrate how dynamic interactions vary one’s sense of agency. From neuroimaging data, we identify regions that are more active when self and other are distinct, including the TPJ. Conversely, we observe activity in the cerebellum, EBA and SMA when self and other blur. These findings suggest that a certain degree of self-other distinction at sensorimotor, experiential, and neurophysiological levels is required to maintain successful interpersonal coordination.

## Introduction

Interacting with the world and with others often requires the attribution of agency: assessing who was responsible for generating an action and its effects. In previous accounts, the definition of agency and its neural correlates has been based primarily on passive and often static tasks of agent attribution. A shift towards a more enactive concept of agency has prompted recent work to describe a more flexible, dynamic and interactive description of self (Fuchs and de Jaegher, 2009; Nahab et al., 2011; Synofzik et al., 2008). Using functional magnetic resonance imaging (fMRI) and an interactive simulated joint action task (paced finger tapping), we examined how dynamic coupling (Dumas et al., 2014b) between a human tapper and an adaptive virtual partner (VP) – producing more or less similar tones – affects coordination performance as well as the sense of self- or joint agency. Manipulating VP adaptivity (Fairhurst et al., 2013) and self-related auditory feedback, we tilted the balance of segregation and integration of self and other information to investigate the neural correlates of varying degrees of perceived self-other distinction.

Previous neuroimaging research on agency has employed various methods to probe the sense of self (Sperduti et al., 2011). Some have done so by contrasting it with a *sense of other*, using perspective switching and observation tasks (Ruby and Decety, 2001). Based on a classical sensory comparator feedback/feedforward model (Farrer and Frith, 2002; Wolpert and Flanagan, 2001), others have tested the effects of matched and mismatched expectations (Spengler et al., 2009). The comparator model posits that the action control system, when specifying a sequence of motor commands to reach a certain goal, creates an efferent copy of these commands. The efferent copy is then used by a forward model to generate a prediction about the next state of the system, which is subsequently compared to the system’s actual state. Congruency between predicted and actual states results in agency being attributed to self while a mismatch is attributed to external causation. In an extension of the comparator model, Nahab and colleagues proposed neural mechanisms for *degrees* of perceived sense of self (Nahab et al., 2011). Related work has identified several key brain areas that are pertinent for self-related processing, including those involved in mismatch detection (insula, inferior parietal lobule, dorsolateral prefrontal cortex - dlPFC and temporoparietal junction - TPJ, cerebellum and extrastriate body area - EBA), synchronous or congruent self-generated actions (supplementary motor area - SMA) and intentional binding (pre-supplementary motor area - pre-SMA, (David, 2012; Nahab et al., 2011). However, with only a few exceptions (Guionnet et al., 2011), these approaches have not examined agency within a *social and interactive context* (David, 2012; Schilbach et al., 2013) and do not distinguish between feelings and judgments of agency, that is, implicit and explicit senses of agency (Miele et al., 2011; Synofzik et al., 2008). Most recently, using a temporal coordination task, Bolt & Loehr detail the nature of so-called joint agency in which, under certain circumstances of coordinated joint action, participants feel a sense of shared control (Bolt and Loehr, 2017).

In the present study, we model dynamic joint action in a controlled fashion by implementing an adaptive auditory “virtual” partner (VP) in a sensorimotor synchronization task (Repp and Keller 2008; Fairhurst et al. 2013). Sensorimotor synchronization, which is intrinsically linked to music making and other forms of joint action, is typically studied by requiring an individual to coordinate simple movements, such as finger taps, with an auditory sequence produced by a computer or a co-acting partner (Ivana Konvalinka et al., 2010; Repp, 2005; Repp and Su, 2013). In our setup, participants were told that they would be tapping with a computer-controlled VP which, to simulate various potential human partners, would vary its tone onsets based on the participant’s tapping performance (Fairhurst et al., 2013, 2014; Koehne et al., 2016). These VPs could adapt highly, moderately, or not at all; that is, account and compensate for the full amount of the asynchrony (the time between the human tap and the computer tone), only a small fraction thereof, or simply maintain a rigid and steady tempo like a typical metronome. These three levels of adaptivity were implemented by modulating the amount of temporal error correction employed by the VP, which computed each asynchrony online and adjusted the timing of its next tone in a compensatory fashion based on a proportion of the previous asynchrony. As in our previous studies (Fairhurst et al., 2013; 2014), we expect that varying VP adaptivity will affect synchronization performance and result in a shift in synchronization strategy related to the use of error correction. This may occur due a perceived variance in the perceived reliability of the VP. Based on our previous studies, coupling is said to be optimal when both parties adapt and employ similar, moderate degrees of phase correction (Repp and Keller 2008). When the VP corrects more than the observed optimal, moderate degree, participants may regard the partner as more erratic or unstable, because overall synchronization between VP and the participant is poor despite large corrective adjustments of the VP.

In contrast with our previous studies, in the present experiment, participant’s taps could produce auditory feedback that varied between trials (“off”-no tone produced, “distinct” –tapping produced a different, easily distinguished, conga drum tone; “ambiguous”: the VP and the participant’s tapping produced the same conga tone). Depending on the condition of auditory feedback, we assumed that the nature of the timbral cues would either facilitate the segregation of auditory streams and agency attribution, thus leading to greater self-other distinction (“distinct” feedback), or, in the case of the same tones, result in self-other blurring (“ambiguous” feedback). In other words, when feedback tones are different for the participant and the VP, stream segregation and agency attribution should be relatively easy; when the feedback tones are the same for both the participant and VP, self and other stream integration should be easy and segregation difficult. Mapped to a musical context, in the latter case, one might expect that individuals playing the same instrument with similar timbres may find it more difficult to distinguish themselves acoustically from their fellow players due to acoustic masking (Meyer, 2009).

Across manipulations of VP adaptivity and self-auditory feedback, the participants’ task was to synchronize their taps with the tones of the VP and to maintain a steady tempo while doing so. The goal of achieving stable synchronization requires the participant’s action control system to determine whether each self-produced tap was early or late relative to the VP tone to compensate for the asynchrony by implementing temporal error correction in the correct direction (lengthening the next inter-tap interval for early taps and shortening the next inter-tap interval for late taps). Under conditions in which the participant’s taps produce tones, the action control system must identify the source of the perceived tones (self vs. other) and determine their temporal order to implement error correction effectively (e.g., if the self-produced tone is earlier than the other-produced tone, then the next inter-tap interval must be lengthened to produce the upcoming self-produced tone slightly later). Determining the temporal order of self- and other-produced tones may be achieved more reliably or efficiently when auditory feedback from the participant and VP are distinct than when feedback is ambiguous. Accordingly, the participant’s use of error correction may be more effective—hence sensorimotor synchronization most stable—with distinctive auditory feedback, and this effect may be most pronounced when VP adaptivity is high. Based on previous work by our group and others, we hypothesized that performance, that is, the stability of synchronization (lower SD asynchrony - a measure indicative of greater stability of the interaction), should vary as a function of feedback type and VP adaptivity. Without auditory feedback, high VP adaptivity leads to instability due to overcorrection (due to additive effects of VP and human error correction, (Elliott et al., 2016; Honisch et al., 2016; Repp and Keller, 2008; Wing et al., 2014) while with distinctive auditory feedback, high adaptivity stabilizes performance (because the comparator can use this information effectively, and the human can dampen error correction); this benefit does not occur with ambiguous feedback because the comparator is comparator is unable to distinguish input related to self or other, and forced to ‘ignore’ the feedback.

After tapping with the VP, participants were prompted to rate task difficulty as well as their sense of influence and sense of oneness in order to gauge judgment of self- or joint agency (Figure 1A). Oneness is used here to describe “the degree of perception of parts of the self in the other” or “the extent of informal or formal tie[s] or the extent of interdependence” (Aron et al., 1992; Cialdini et al., 1997; Davis et al., 1996). This may be linked to similar concepts such as “we-ness” as described by Gallotti and Frith (Gallotti and Frith, 2013) or “togetherness” (Hart et al., 2014), shared or joint agency (Bolt and Loehr, 2017) and in music, this sense of oneness is an aspect of being “in the groove” (Janata et al., 2012). This is relevant to group music making, where not only are movements synchronized but also the produced sounds overlap in time and space, potentially underlying shared representations of self and other (Sebanz et al., 2006). The obtained ratings of oneness serve not only to explore the neural correlates of self-other merging but also to test the hypothesis that to successfully perform the task, that is, maintain a stable level of synchronisation, an element of self-other distinction is also required (Gallotti et al., 2017; Novembre et al., 2016, 2012). We predicted that ratings of oneness would be lowest when tapping with an unstable partner with ambiguous auditory feedback because in this case the variable timing of the VP works against self-other integration while the timbrally indistinguishable tones produced by the participant and VP work against self-other segregation (Bolt and Loehr, 2017).

**Figure 1:**
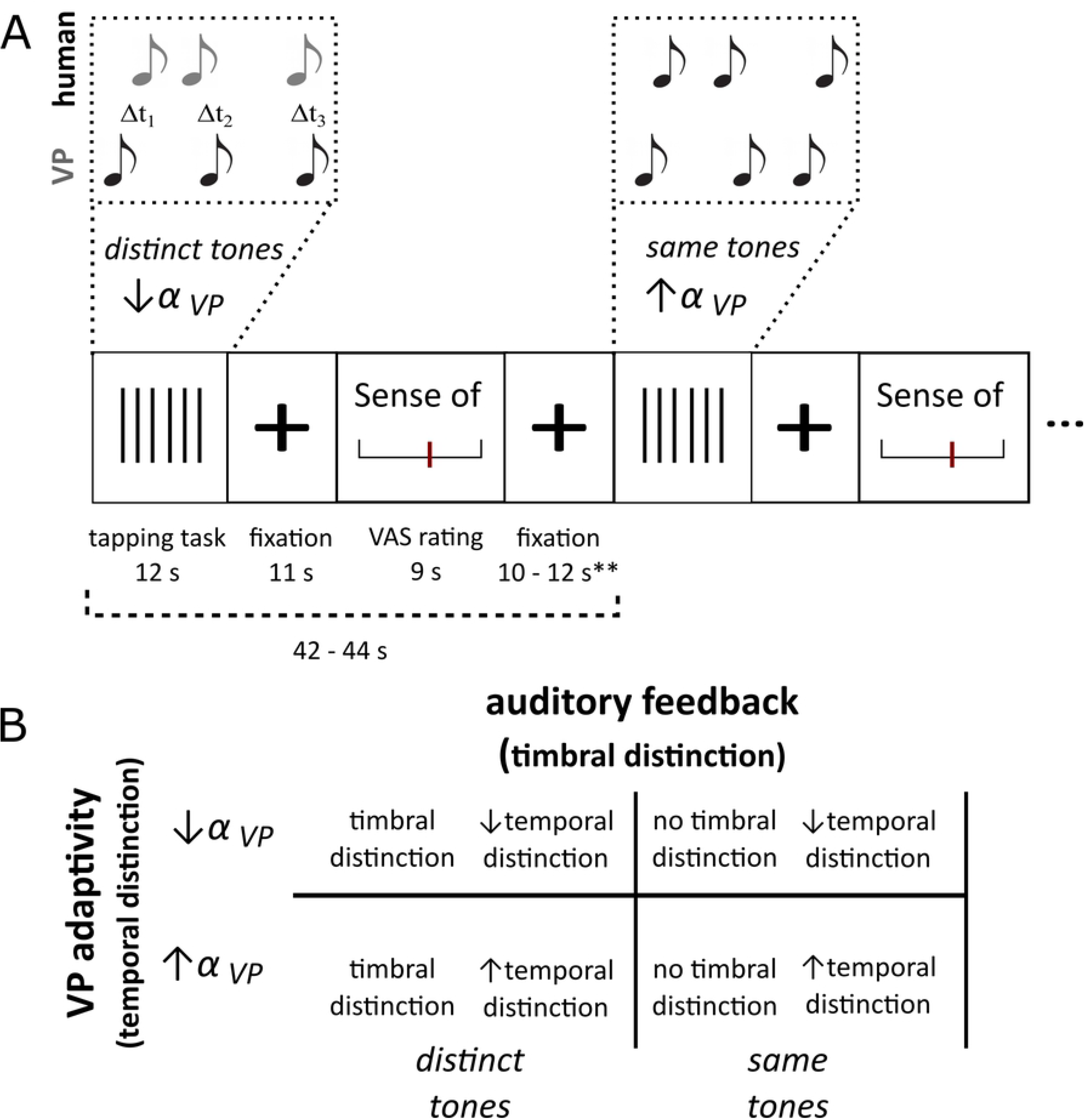
Study design for exploring variable and hierarchical feelings of self-agency. A) fMRI event-related design illustrating timing of tapping block followed by visual anologue scale ratings (for full details, please refer to Methods). B) Model for a two-factor manipulation in which timbral (tone type - auditory feedback) and temporal (virtual partner adaptivity or α_VP_ - stability of VP tone onset) cues result in greater degrees of self-other overlap or distinction as measured by tapping performance (as well as subjective reports of perceived influence (self-agency) and oneness (self-other overlap).

We further hypothesize that manipulation of our two factors (timing and timbre) will affect the perceived influence that the participant feels s/he is exerting within the partnership over the tempo - an assessment that we posit requires an increased awareness of self/other distinction that can be linked to greater self-agency. By obtaining subjective trial-by-trial ratings of perceived influence, we examine a more nuanced (non-binary) relationship between tapping performance and perceived influence predicting that when auditory feedback is distinct, temporal and timbral cues will encourage self-other segregation (Kühn et al., 2013). By contrast, lower ratings of influence (in conjunction with lower perceived oneness) should be observed when tapping with an unstable partner with ambiguous auditory feedback. Using a novel, interactive and dynamic model of temporal coordination which results in varying degrees of self-other overlap (in time and timbral quality), we explore how this affects objective tapping behaviour, subjective experience, and correlated neural activity.

## Materials and methods

### Participants

The experiments were conducted at the Max Planck Institute for Human Cognitive and Brain Sciences in Leipzig, Germany. 16 healthy volunteers (eight females and eight males; age range: 21-33; mean age: 26.31 years, SD=±4.71) were recruited after screening for absence of any prior neurological or psychiatric disorders and contraindications for MR experimentation. Subjects all had previous finger tapping task experience. Written informed consent was obtained for each subject before scanning. The research protocol was in accordance with the Declaration of Helsinki.

### Study Design

During the pre-scan instructions, participants were told that they would be tapping with a VP which would vary its tone onsets based on their tapping performance. Furthermore, it was explained that their taps would produce auditory feedback that varied between trials. The manipulations chosen included varying both the type of auditory feedback (off, on-ambiguous, and on-distinct) and degree of VP adaptivity (non-adaptive, moderately adaptive, and highly adaptive). This 3 × 3 design allowed for four repetitions of each condition within a run, and thus eight repetitions were presented across two consecutively acquired runs. Each imaging session lasted approximately 1 hour and consisted of 36 pseudo-randomized task trials (jittered to last between 42 – 44 seconds) with no cue as to the trial type prior to the start of a trial. Each trial was followed by a rest baseline period of between 10 – 12 seconds. Total scan time for each participant was approximately 40 minutes. (For further details of study design, please refer to Figure 1). For each of the tapping trials, participants were instructed to attend to two isochronous initiation tones which demonstrated the expected tempo and were instructed to synchronize their taps with the tones, starting with the third tone, as accurately as possible and to maintain the initial tempo to the best of their ability. The dual aspects of the instructions—to synchronize *and* to maintain the initial tempo—were intended to help ensure that participants would not be put off by the variability of the VP in the high adaptivity condition.

In each case, starting with the first initiation tone, the fixation cross turned from black to green and was displayed for the duration of the tapping task trial. After the 22 tone-tap pairs of the trial, subjects were cued to provide subjective visual analogue scale (VAS) ratings for the preceding tapping trial using a two-button response box. VAS were presented to obtain online ratings for a randomly chosen pairing of either: sense of *oneness*, *influence*, and *difficulty*.

### Stimuli

#### Auditory Stimuli

The VP was programmed so that, if the participant’s tap preceded the tone, a negative asynchrony was registered and the next sequence inter onset interval (IOI) was shortened (i.e., the next VP tone occurred sooner). Conversely, if the participant’s tap occurred after the tone, a positive asynchrony was registered resulting in a lengthening of the next IOI. The direction of the temporal error correction implemented by the VP was the opposite of the correction expected in the participant’s taps, as it should be if the computer (controlling the tones) ‘‘cooperates” with the participant (controlling the taps). The amount of the correction (α) was varied by a fraction of the calculated asynchrony across three conditions, ranging from no error correction (α=0), moderate error correction (α=0.25, see Fairhurst et al., 2013) to full correction (α=1), in the non-adaptive, moderately adaptive, and highly adaptive conditions, respectively (Fairhurst et al., 2013; Fairhurst et al., 2014). Further details on the algorithm governing the VP are given by Repp and Keller (Repp and Keller, 2008)). The tones were specified to be 50 ms in duration played as synthesized “conga drum” sounds. In the auditory feedback “off” conditions, no tone was produced by participant taps, while in the “on” conditions, participant taps either produced a different drum sound (“distinct”; *mute* and *open hi conga*) to that of the VP or the same conga sound as the VP (“ambiguous”; both *open hi conga*). Participants listened over Siemens MR compatible headphones at a comfortable intensity.

#### Visual stimuli

Visual analogue scales (VAS) were presented to obtain online ratings for a randomly chosen pairing of either: sense of *oneness*, *influence*, and *difficulty* of the preceding trial. For oneness, participants estimated both two estimates a) How in synch they were with the metronome and b) how overlapping the tone of their tap with that of the metronome. The oneness (“Gleichklang”) rating scale was anchored with no sense of oneness (“kein”) at the minimum and with complete sense of oneness (“komplett”) at the maximum. Participants were instructed to rate this bearing in mind both temporal and timbral overlap between their tapping and the tones produced by the VP. We therefore interpret oneness as a subjective measure of perceived self-other merging in time and in the auditory space (tones overlapped). Additionally, we acquired ratings of influence, an assessment which we posit requires an increased awareness of self as distinct from other. Participants were instructed to rate how much their tapping influenced the overall rhythm using a scale “Influence” (“Einfluss”) scale anchored with no influence (“kein”) at the minimum and absolute influence (“absolute”) at the maximum. The rating of influence was described to participants as a measure of how much they felt their tapping performance influenced the overall tempo. Similarly, the “Difficulty” (“Schweirigkeit”) scale was anchored with very easy (“sehr leicht”) and extremely difficult (“sehr schweirig”) and was explained as a measure of how difficult it was to synchronise with the virtual partner. Ratings of influence and oneness were sampled three times per condition. Difficulty ratings were acquired to control for potential effects of difficulty due to the manipulation (sampled twice per condition). Each scale was presented for 4.5 seconds. All visual stimuli were projected onto a screen visible to the subject via prism glasses. Visual stimulation was thus continuous throughout the experiment.

### Data acquisition

#### SMS Tapping Data acquisition

Participants were instructed and trained to tap with their right index finger on an in-house built, MR-compatible air-pressure tapping pad that was connected to the computer via a MIDI interface. Taps were recorded by MAX 4.5.7 http://www.cycling74.com).

#### MRI Data acquisition

Functional imaging was conducted using a 3 Tesla Siemens Trio system. An echo-planar imaging (EPI) sequence was used with the following pulse sequence parameters: TR = 2000 ms; TE = 24 ms; 36 × 3 mm axial oblique slices; 1 mm gap; voxel size = 3 × 3 × 3 mm^3^; volumes = 699. Scans were acquired continuously throughout the experiment. High resolution, T1-weighted, structural scans (64 slices at 1 × 1 × 1 mm^3^ voxel size) were obtained for each individual for anatomical overlay of brain activation.

### Data Analysis

#### Subjective Ratings

Individual means, grouped by type of auditory feedback and degree of VP adaptivity, were calculated for the ratings of perceived oneness, influence over the pulse, and difficulty to synchronize during tapping tasks. To do so, VAS ratings were converted into numerical 0-10 ratings. A group mean and standard deviation was calculated for the post scan overall subjective rating of performance to be compared with objective measures of synchronized tapping. Individual mean ratings were used to compare with the objective tapping data and were subject to standard ANOVA tests to probe main and interaction effects of the two manipulated factors (auditory feedback, VP adaptivity).

#### Sensorimotor Synchronization Data

The recorded times of computer tones and human taps were analyzed in terms of asynchronies, defined as the differences between tone and tap onsets. The standard deviation (SD) of asynchronies provides an index of the stability of sensorimotor synchronization performance, such that smaller SDs of asynchronies indicate greater stability in sensorimotor synchronization. We assume that SD asynchronies decreases as self–other focus increases due to increased coupling strength between the human participant and the VP. Moreover, we investigated coupling strength, as reflected by the degree of error correction implemented by the human participant (α_human_) and which represents the degree to which the human adapted to asynchronies (i.e., the average proportion of each asynchrony that is compensated for by adjusting the time of the next tap). It can be estimated by calculating the zero crossing point of lag-1 autocorrelation functions of the asynchronies across VP adaptivity conditions (0, 0.25 and 1) (Repp & Keller, 2008). These tapping measures were calculated both within and across subjects, across conditions of VP adaptivity and used here to further explore the fMRI data.

#### Imaging Analysis

Analysis of all neuroimaging data sets was performed using FEAT (FMRIB Expert Analysis Tool) Version 5.63, part of FSL (FMRIB’s Software Library, www.fmrib.ox.ac.uk/fsl). Pre-statistic processing included: motion correction using MCFLIRT (Motion Correction FMRIB’s Linear Image Registration tool, (Jenkinson, 2001), non-brain removal using BET (Smith, 2002), spatial smoothing using a Gaussian Kernel of 4 mm full width at half-maximum and non-linear high pass temporal filtering (Gaussian-weighted least-squares straight line fitting, with sigma=40.0 s). Registration included co-registration of the functional scan onto the individual T1 high-resolution structural image and then registration onto a standard brain (Montreal Neurological Institute MNI 152 brain) using FLIRT (FMRIB’s Linear Image Registration Tool, (Jenkinson, 2001).

Statistical analysis at the first, individual subject level was carried out using a general linear modeling (GLM) approach (Friston KJ, 1994). Time-series statistical analysis was carried out using FILM (FMRIB’s Improved Linear Model) with local autocorrelation correction (Woolrich MW, 2001). Second level analysis grouped the data of each subject’s two scanning blocks, using the data from the first level of analysis. For group statistics, analysis was carried out using FEAT (FMRI Expert Analysis Tool) with higher-level analysis carried out using FLAME (FMRIB’s Local Analysis of Mixed Effects). This analysis method allows for incorporation of variance within session and across time (fixed effects) and cross session variances (random effects). Cluster thresholding was performed with a Z-threshold of 2.3 and a corrected p-value of < 0.05 with a cluster-based correction for multiple comparisons using Gaussian Random Field Theory (Worsley KJ, 1992; Friston KJ, 1994).

Contrasts were calculated to identify activation during each of the nine conditions compared to baseline. Each task block was modeled as two defined events: initiation (perception of initiation tones) and synchronized tapping. Subtraction contrasts between conditions were also performed. Based on the behavioural data, the design was simplified to include only conditions in which both “self” and “other” auditory feedback was available resulting in a 2×2 comparison with factors of VP adaptivity (moderately adaptive and highly adaptive) and auditory feedback (ambiguous vs. distinct). A 2 × 2 repeated measures ANOVA was run to explore the main and interaction effects of auditory feedback and VP adaptivity within the four conditions of interest (moderately adaptive distinct, moderately adaptive ambiguous, highly adaptive distinct, highly adaptive ambiguous). Individual means of perceived difficulty ratings per condition were used to control for experienced task difficulty.

Regression analyses were conducted identifying areas of covariance between BOLD signal change during tapping with factors of subjective and objective task performance as well as subjectively perceived degrees of influence and oneness. This was done by incorporating individual means per condition and entering these as separate regressors. Functionally and individually defined region masks were created to perform exploratory region of interest (ROI) analyses to further describe activity across conditions. Masks for the anterior insula, EBA, posterior cerebellum and SMA were based on the subtraction contrast from the “ambiguous”: highly adaptive vs. moderately adaptive conditions, areas that have previously been identified as relevant for processing of agency and agency violation (Yomogida et al., 2010). Additionally, a mask for right TPJ was created based neural activation from the 2×2 ANOVA, main effect of auditory feedback.

## Results

### Tapping behaviour

In the objective tapping behavior data, there was a significant interaction of our two factors (3 × 3 ANOVA: no significant main effects, significant interaction: F(2,56)=5.147, p<.05 – see Table 1). Specifically, in the highly adaptive VP condition (α=1) which otherwise would result in poorer synchronization, individuals were significantly more stable (relative to the “off” condition) when they tapped in the distinct compared with the ambiguous auditory feedback condition (distinct vs. off: t(15)=2.76 p<.05; ambiguous vs. off: t(15)=0.279 p=.784 – see Table 1).

**Table 1:**
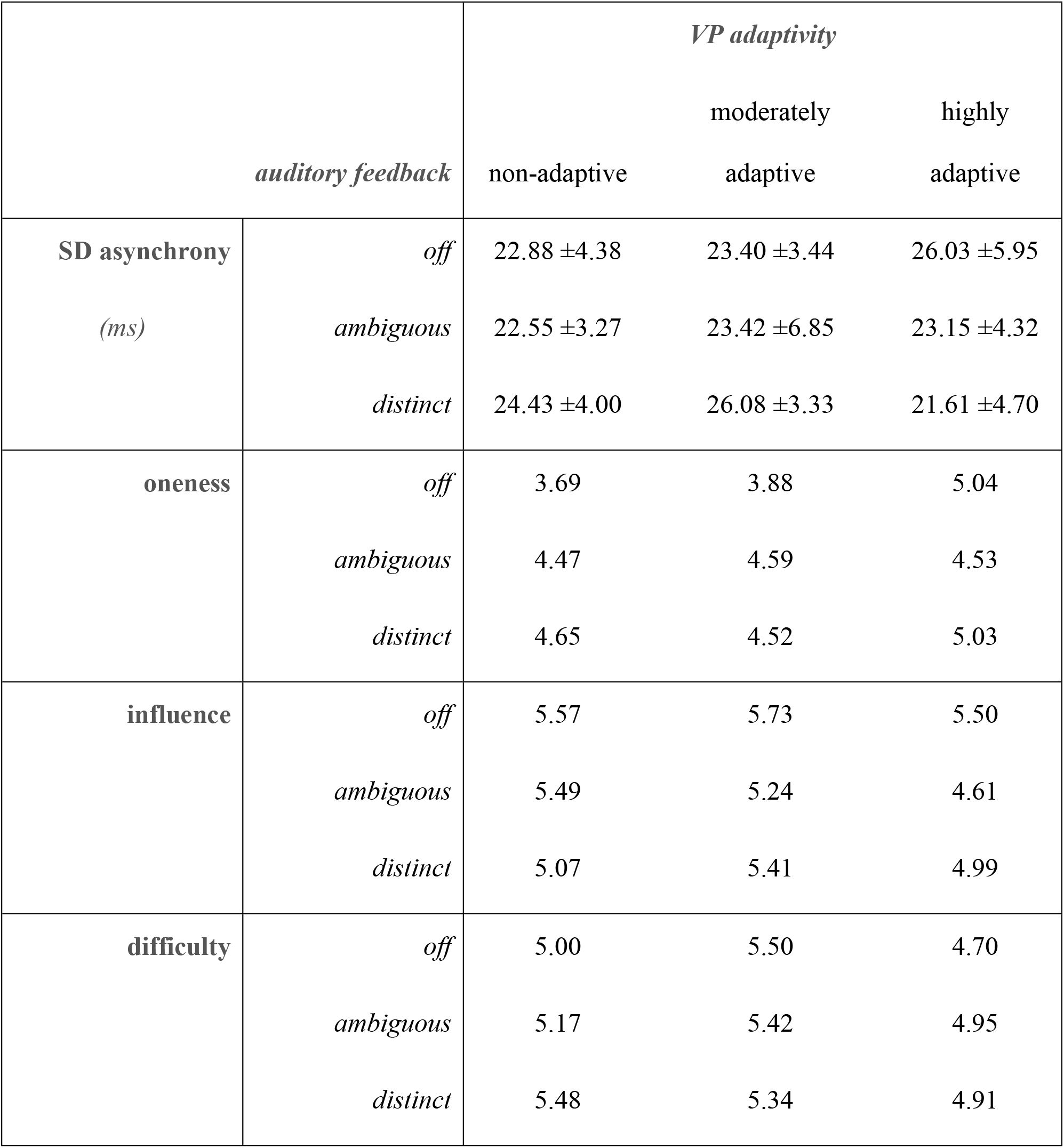
Behavioural data. Group mean behavioural data of tapping performance as measured by standard deviation of asynchronies (SD asynchrony, group mean ± standard deviation) and subjective ratings of perceived oneness, influence and task difficulty (group mean).

|  |  | *VP adaptivity* |  |  |
| --- | --- | --- | --- | --- |
|  |  | non-adaptive | moderately<br>adaptive | highly<br>adaptive |
| **SD asynchrony**<br>(*ms*) | *off* | 22.88 ± 4.38 | 23.40 ± 3.44 | 26.03 ± 5.95 |
| | *ambiguous* | 22.55 ± 3.27 | 23.42 ± 6.85 | 23.15 ± 4.32 |
| | *distinct* | 24.43 ± 4.00 | 26.08 ± 3.33 | 21.61 ± 4.70 |
| **oneness** | *off* | 3.69 | 3.88 | 5.04 |
|  | *ambiguous* | 4.47 | 4.59 | 4.53 |
|  | *distinct* | 4.65 | 4.52 | 5.03 |
| **influence** | *off* | 5.57 | 5.73 | 5.50 |
|  | *ambiguous* | 5.49 | 5.24 | 4.61 |
|  | *distinct* | 5.07 | 5.41 | 4.99 |
| **difficulty** | *off* | 5.00 | 5.50 | 4.70 |
|  | *ambiguous* | 5.17 | 5.42 | 4.95 |
|  | *distinct* | 5.48 | 5.34 | 4.91 |

To further investigate how the different types of auditory feedback affected the underlying sensorimotor dynamics between the human tapper and the VP, we estimated the degree of human error correction (mean ±SD: *off* 0.37±0.14, *distinct* 0.29±0.12, *ambiguous* 0.37±0.13). Our data show a significant effect of human error correction (f(1.746,26.185)=4.378, p<.05) with participants employing the least error correction in the “distinct” condition (*off vs. distinct:* t(15)=2.54 p<.05, *distinct vs. ambiguous:* t(15)=−2.73 p<.05 – see Table 1).

### Subjective measures of perceived oneness and influence

Oneness ratings varied as a function of both degree of VP adaptivity as well as the distinguishability of self and other tones (main effect of auditory feedback F(2,60)=6.90, p<0.01; main effect of VP adaptivity F(2,60)=3.44, p<0.05). Overlap in tone type (“ambiguous” timbral cues) resulted in greater perceived oneness but only when tapping with a reliable partner (moderately adaptive VP with ambiguous feedback vs. no feedback: t(15)=3.01, p<0.01); highly adaptive VP with ambiguous feedback vs. no feedback: t(15)=−0.66, p=0.52). When tapping with an unstable (highly adaptive) partner, oneness ratings were greater when distinguishable self-other information was available (“distinct” timbral cues). Tapping with an ambiguous and highly adaptive partner resulted in lower ratings of perceived oneness (highly adaptive VP – distinct vs. ambiguous feedback: t(15)=2.12 p<0.05). Additionally, we observed a main effect of VP adaptivity on perceived influence ratings (F(2,60)=3.91, p<0.05; no significant main effect of auditory feedback or interaction) with a significant decrease in the degree of perceived influence with the most unstable, highly adaptive VP (Table 1). Difficulty ratings confirmed that across conditions, participants were equally challenged (no main effect or interaction of VP adaptivity or auditory feedback on difficulty ratings – see Table 1). This suggests that differences influence or oneness are not attributable to task difficulty.

### Neural correlates of self-other distinction and merging

Based on the pattern of behavioral data, we explored the imaging data as a reduced 2 (auditory feedback: distinct versus ambiguous) × 2 (VP adaptivity: moderately adaptive versus highly adaptive) factorial design, i.e. including only instances in which both “self” and “other” auditory feedback were available.). In effect, the participants can be seen as having interacted with one of four possible (virtual) partners: i) a moderately adaptive, fairly stable but distinctly sounding “other”, ii) a moderately adaptive, fairly stable but ambiguously sounding “other”, iii) a highly adaptive, unstable but distinctly sounding “other” or iv) a highly adaptive, unstable but ambiguously sounding “other”. A two-way ANOVA showed a significant main effect of auditory feedback and a significant interaction between auditory feedback and level of VP adaptivity. The main effect of auditory feedback, showing variance associated with either distinct or ambiguous self-related information, correlated with bilateral activation of the temporoparietal junction (TPJ) and auditory cortex (planum temporale) (Figure 2A, Table 2A) with greater activity with distinct timbral cues, marked behaviourally with individuals correcting for a smaller proportion of each asynchrony (compared with the ambiguous condition, see Figure 2B).

**Table 2:** 2×2 ANOVA Auditory feedback × VP Adaptivity (repeated measures ANOVA). Coordinates in MNI space and associated peak voxel Z-scores. p < 0.05 corrected for multiple comparisons.

| Main effect of auditory feedback |  |  | Peak MNI |  |  |
| --- | --- | --- | --- | --- | --- |
|  |  |  | Coordinates |  |  |
| Regions |  | Z-max | x | y | z |
| Planum temporale | R | 7.42 | 56 | -14 | 4 |
| Planum temporale | L | 7.42 | -48 | -14 | 4 |
| Temporoparietal junction | R | 4.62 | 58 | -40 | 8 |
| Temporoparietal junction | L | 3.58 | -50 | -58 | 16 |
| **Interaction (auditory feedback x VP adaptivity)** |  |  |  |  |  |
| Anterior insula | L | 4.32 | -40 | 16 | -2 |

**Figure 2:**
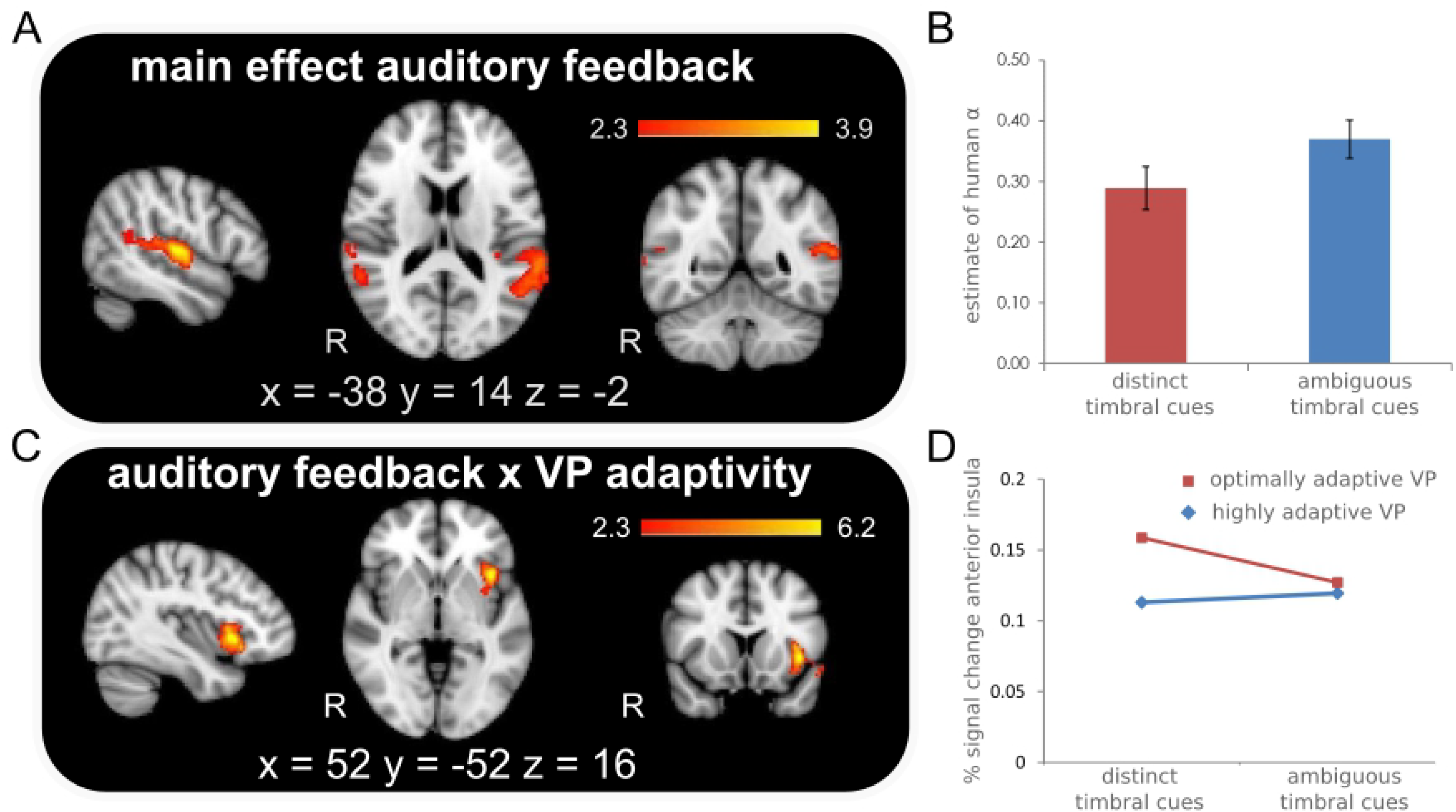
Effects of auditory feedback and VP adaptivity. Imaging results from 2×2 repeated measures ANOVA (random effects, Z = 2.3; p = 0.05, corrected), where A) shows main effect of auditory feedback with activity in planum temporale and temporoparietal junction, B) shows group average estimate of error correction (α) across distinct and ambiguous auditory feedback conditions, C) shows interaction of auditory feedback and VP adaptivity with isolated activation of anterior insula and D) shows results from an ROI analysis of the left anterior insula across conditions of VP adaptivity (optimally adaptive vs. completely adaptive VP) and across conditions of auditory feedback (distinct and ambiguous auditory feedback). See Table 2 for full list of activation with MNI coordinates.

Additionally, a significant interaction between our two manipulated factors was isolated to the left anterior insula (Figure 2C, Table 2B). An ROI analysis of the anterior insula across timbral cue types (distinct vs. ambiguous), comparing conditions of tapping with either a moderately adaptive partner or a highly adaptive partner revealed greater activity in the highly adaptive condition when timbral cues are distinct (“distinct”) but no difference in activation between VPs in the “ambiguous” condition (Figure 2D). This suggests a greater sensitivity of the anterior insula to discriminating timbral cues when dealing with a more unreliable partner. Probing the direction of the main effect further, we contrasted the conditions in which individuals tapped with a moderately adaptive partner with either distinct or ambiguous auditory feedback. When the temporal cues were more reliable and the timbres were distinct, a paired sample t-test showed greater activation in bilateral auditory cortex and TPJ and additional activity in the right ventrolateral prefrontal cortex (vlPFC) (Figure 3A, Table S1A). Additionally, an ROI analysis of the right TPJ across all levels of VP adaptivity (i.e. irrespective of the reliability of the partner and thus specific timing cues) showed that activity in this structure was greatest during the distinct condition compared to the ambiguous auditory feedback condition. Similarly, an ROI analysis of the SMA showed a significant difference between distinct and ambiguous conditions with greater SMA activity in response to distinguishable self-related auditory information.

**Figure 3:**
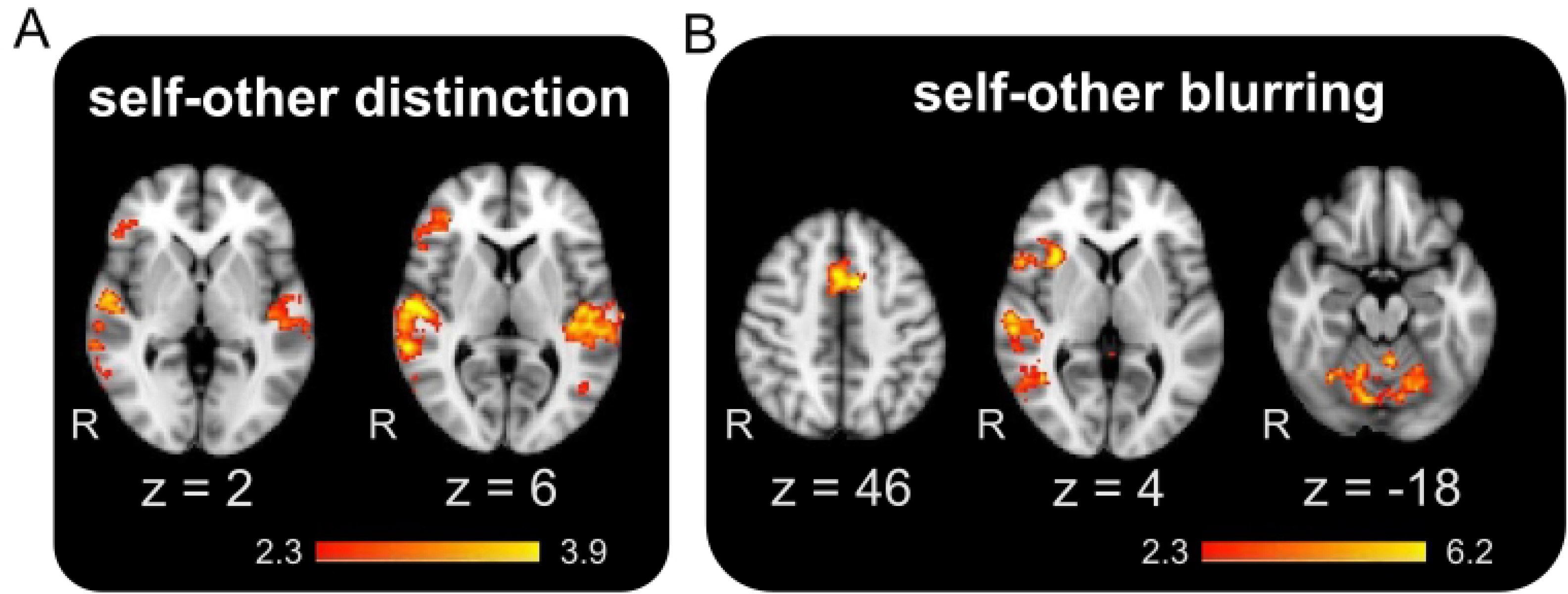
Self-other distinction and blurring. Imaging results for A) paired sample t-test contrast of distinct vs. ambiguous auditory feedback when tapping with a moderately adaptive VP (random effects, Z = 2.3; p = 0.05, corrected), B) paired sample t-test contrast of tapping with a highly adaptive VP vs. optimally-adaptive VP while receiving ambiguous auditory feedback (random effects, Z = 2.3; p = 0.05, corrected). See Table S1 for full list of activation with MNI coordinates.

In our behavioral data, we saw that at high levels of VP adaptivity, the erratic nature of the partner in conjunction with ambiguous auditory feedback resulted in decreases of both perceived oneness and influence. A subtraction contrast of conditions in which “ambiguous” feedback was given tapping with either highly adaptive VP or an optimally adaptive VP showed areas previously suggested to be involved in agency violation including the cerebellum, SMA, EBA, IFG and anterior insula to be more active when tapping with a highly adaptive partner (Figure 3B, Table S1B). A regression analysis incorporating individual trial-by-trial ratings of oneness was performed identifying positively correlated activation with increasing degrees of perceived oneness in the precuneus, inferior parietal lobule and posterior cingulate (Table S1C).

## Discussion

Our interactive finger-tapping task had participants synchronize their taps with the tones of a computer-controlled VP that was programmed to adapt to participants’ tap timing to varying degrees. Better task performance, that is stable synchronization, required participants to identify the source of the created tones (self and other) and then calculate and compensate accordingly for the asynchrony. Our manipulation varied (1) VP adaptivity and therefore the variability in the onset of the VP’s tones, and (2) the quality of the self-related auditory feedback. As such, the task induced variation in available temporal and timbral cues and the resulting sense of self-agency, assessed by ratings of perceived influence (indexing self-other distinction) and oneness (self-other merging). Moreover, our design distinguishes between a subconscious level processing of agency that underlies the task and the explicit judgments of agency in the form of our ratings.

We showed that the availability of additional sensory information pertaining to the self (beyond proprioceptive feedback) affected performance and our measures of self-other distinction (influence) and of self-other merging (oneness). Together, these results provide evidence for the evolving concept of a variable sense of self-agency and highlight the importance of relevant activity in brain areas associated with agency attribution, agency violation and self-other distinction. Specifically, we identified regions that were more active in conditions in which self and other were distinct, including the TPJ. Conversely, increasing degrees of perceived oneness were seen to correlate with activity in the precuneus, inferior parietal lobule and posterior cingulate. Additionally, we observed activity in the cerebellum, EBA and SMA when self and other blurred. Both our behavioral and neural results suggest that while some degree of self-other merging may be useful for and result from successful joint-action, a certain level of self-other distinction is required to maintain a sense of self agency (Keller, P.E., Novembre, G., & Loehr, 2016; Novembre et al., 2016, 2012).

### The importance of self-related information when tapping with an adaptive virtual “other”

No significant main effects of either auditory feedback or VP adaptivity were seen on our tapping measure of synchronization stability suggesting that any effects on our dependent variables (behavioral or neural) were not due to differences in task performance. We did, however, see a significant interaction of our manipulated factors with distinct self-related information resulting in more stable synchronization in the highly adaptive VP condition. This improvement in performance corroborates and extends our understanding of the effect of additional auditory feedback on sensorimotor synchronization (Aschersleben and Prinz, 1995). We explain the observed improvement in synchronization stability in the distinct, highly adaptive VP condition by the fact that when the signal from the “other” agent is less reliable, distinct self-related information is prioritized and most useful (I. Konvalinka et al., 2010). Moreover, this finding complements recent work that explored how task performance and agency varied in continuous action tasks (Inoue et al., 2017; Wen et al., 2015).

Additionally, we saw further evidence of this shift in prioritization with calculated estimates of error correction employed by the participants. We found that in the ambiguous and “off” conditions, individuals corrected for a larger proportion of each asynchrony than in the distinct condition. At a neural level, this shift in focus from self-other to self, linked to the significant interaction of our factors may explain activity of the left anterior insula. The anterior insula forms part of a spatial attention network (Mayer et al., 2006) and has also been shown to be activated in tasks involving time perception (Livesey et al., 2007). Furthermore, this area is frequently associated with tasks which involve self-awareness (Craig, 2009) and has more specifically been linked to self-reflection (Modinos et al., 2009). Interestingly, oneness, defined to participants as sounding alike and tapping together with the partner, was greatest when tapping with a distinctive, highly adaptive partner, suggesting that oneness is dependent on a certain level of self-awareness (not on producing overlapping sounds).

A large meta-analysis revealed that anterior insula activity is commonly seen in tasks involving self, as opposed to externally, attributed agency (Sperduti et al., 2011). Though some reports associate the right anterior insula with self-related processing (Uddin, L. Q., Iacoboni, M., Lange, C., & Keenan, 2007), our thresholded activation was seen unilaterally on the left (Modinos et al., 2009). The precise nature of the functional lateralization of this area is as yet unclear. More recently, left lateralized activity has been associated with cognitive control processes important for subsequent behavioral adaptations (Späti, J., Chumbley, J., Brakowski, J., Dörig, 2014). Consistent with this, our ROI analysis revealed that activity in the anterior insula varied as a function of the importance of, or reliance on, the timbral cues, with greater activity when tapping with a highly adaptive VP when distinct self-other tones were produced.

Beyond highlighting the fact that the degree of self-focus and thus the importance of self-related information can vary, our manipulation of VP adaptivity is also useful for delineating and describing the difference between objectively having more control over the overall tempo (as the VP compensates either moderately or fully for asynchronies), and subjective ratings of perceived influence. The present findings bring to light the idea of a hierarchy of agency processes: lower (e.g. sensorimotor as indexed by variation in our measure of error correction) and higher (e.g. explicit judgements of agency in the form of subjective ratings) (David et al., 2008). Moreover, previous work by Nahab and colleagues have proposed the idea of a relative and dynamic sense of influence and agency (Nahab et al., 2011). In our case, perceived influence, as a proxy for self-other distinction, varies depending on the nature of the partner and similarity between self and other. Specifically, in the highly adaptive condition, when individuals objectively had the greatest sway within the partnership but received ambiguous self-related feedback, ratings of influence were lower.

### Self-other distinction and self-other blurring

The main effect of auditory feedback identified isolated bilateral posterior activation of the planum temporale and TPJ. These areas of the so-called spatial, or “where”, pathway are commonly activated by tasks involving auditory scene analysis (Alain et al., 2001). An ROI analysis exploring activity in the TPJ across conditions of auditory feedback showed that activity was greatest in the distinct condition. This may be the result of greater segregation in the “distinct” auditory condition in which self and other tones are more easily identified and streamed. Moreover, related work has identified activation of the planum temporale in tasks in which the temporal relationship between parts is used as a cue for stream segregation (Ragert, M., Fairhurst, M. T., & Keller, 2014). Our behavioural tapping data revealed that in the “distinct” condition participants were adapting less to the virtual partner (lower human error correction), which may reflect less integration. To describe the direction of the main effect of auditory feedback, we contrasted the distinct vs. ambiguous conditions (at a moderate level of VP adaptivity) again identifying greater activity in the distinct condition in planum temporale and TPJ activity as well as SMA and right IFG, structures that collectively are implicated in agency processing. Kühn and colleagues suggest that activity in the TPJ is related to matching action with its effects (Kühn et al., 2013). Additionally, observed TPJ activity may be related to several aspects of self-processing, such as agency, self–other distinction, switching between self and other representations and mental own-body imagery (Blanke and Arzy, 2005; Ruby and Decety, 2001; Spengler et al., 2009; Sperduti et al., 2011; Vogeley and Fink, 2003). Similarly, looking at neural overlap and differences between action monitoring and judgments of agency, Miele and colleagues identified TPJ activity which they attributed to either monitoring the comparator signal or for executing corresponding motor corrections (Miele et al., 2011).

Our manipulation also allowed for instances in which there would be greater likelihood of objective (temporal synchronization) and perceived self-other (produced tones) overlap. Behaviorally, we included a rating of oneness to investigate the nature of the perceived self-other merging. Our data identify a relationship between perceived oneness and the degree of self-agency, with lower ratings of oneness in conditions in which influence was rated as low. Where do my actions end and yours begin? One may become confused in social interactions in which one’s movements are highly coordinated with those of others. In group music making, this is almost certainly the case as not only are movements synchronized but the resulting sounds produced can overlap in time and space (Keller, 2014). It has been suggested that it is only once agency is lost that it becomes salient and conscious (Kühn et al., 2013). In the present study, ratings of oneness and influence were lower when tapping with an unstable partner in conjunction with ambiguous auditory feedback. Work by Bolt and Loehr has described a relationship not only between an individual’s own performance stability and joint agency but also between reliability of a co-actor’s actions and a shared sense of control (Bolt and Loehr, 2017). Previous work has suggested that assessing and attributing agency is dependent on the sensory reliability and distinctiveness of the signal (Elliott, M. T., Wing, A. M., & Welchman, 2014). This issue in particularly germane in choral singing, for example, where individuals adjust the intensity of their vocal output in order to optimize the so-called “self-to-other ratio”, which reflects the degree to which an individual can hear their own sounds amongst co-performers’ sounds (Ternström, 2003).

In our neural data, we observed that tapping with the unstable, ambiguous partner resulted in activity in areas previously implicated in agency error or violation including the cerebellum, SMA, EBA, IFG and anterior insula (Yomogida et al., 2010). Activation of EBA in conjunction with these sensorimotor areas may also provide evidence for a network of areas involved in central monitoring or a comparator mechanism underlying agency. Though in separate observations, the EBA and TPJ have both been linked to processing of agency violation and mismatch detection (Sperduti et al., 2011; Yomogida et al., 2010).

### Conclusions

Self-agency has been suggested previously to depend on and be affected by interactions with one’s environment, though the relative contribution of resultant sensory and timing cues for agency may vary (Knoblich and Repp, 2009; Rohde and Ernst, 2016). Importantly, this reliance is thought to depend on action and movement. Recent debate in the field of social cognition has highlighted the need for more active, participatory approaches to describe joint action and agency (Schilbach et al., 2013). In the current study, we propose a dynamic experimental approach to investigate self-agency during interpersonal cooperation (Dumas et al., 2014a; Kelso et al., 2009; Tognoli, E., & Kelso, 2015). Both behavioural and neural data highlight the need for both self-other merging and self-other distinction in temporal coordination tasks, reinforcing the idea that the concept of agency should not be seen as a “static dichotomy … but rather a gradual and highly plastic process that allows the subject to constantly redefine the causal relations to its surroundings” (Synofzik et al., 2006). Understanding the dynamics of this process is as challenge for future research on real-time interpersonal coordination.

